# Protective low avidity anti-tumour CD8+ T cells are selectively attenuated by regulatory T cells

**DOI:** 10.1101/481515

**Authors:** G. Sugiyarto, D. Prossor, O. Dadas, T. Elliott, E. James

**Author notes:** Correspondence: Edd James Centre for Cancer Immunology, University Hospital Southampton, Tremona Road, Southampton SO16 6YD.

## Abstract

Regulatory T cells (Treg) play a major role in the suppression of protective anti-tumour T cell responses. In the CT26 BALB/c murine model of colorectal carcinoma, Tregs differentially suppress responses to two characterised CD8+ T epitopes, AH1 and GSW11, which results in an absence of detectable IFN-γ producing GSW11- specific T cells in the spleen and lymph nodes of tumour challenged mice. Activation of GSW11-specific T cells correlates with protection against tumour growth. Here we show that GSW11-specific T cells are in fact induced in Treg-replete, CT26-bearing mice, where they make up the majority of tumour infiltrating CD8+ lymphocytes, but exhibit a dysfunctional ‘exhausted’ phenotype. This dysfunctional phenotype is induced early in the anti-tumour response in draining lymph nodes, spleens and tumours and is significantly more pronounced in GSW11-specific T cells compared to other tumour-specific T cell responses. Depletion of Tregs prior to tumour challenge significantly reduces the induction of exhaustion in GSW11-specific T cells and correlates with altered T cell receptor (TcR) usage. Moreover, the avidity of GSW11- specific TcRs that expanded in the absence of Tregs was significantly lower compared to TcRs of cytotoxic T lymphocyte (CTL) populations that were diminished in protective anti-tumour responses. This indicates that Tregs suppress the induction of protective anti-tumour T cell responses and may signify that the induction of low avidity T cells, while being more susceptible to exhaustion are the most efficacious in tumour rejection.

## Introduction

CD8+ T cell responses directed to tumours have been shown to occur in many human cancers, where they are a positive prognostic indicator (1-5). Backed by studies in preclinical mouse models, which clearly show that CD8+ T cells are important in the clearance of tumours and may confer lifelong protection against malignancy (6, 7), immunotherapies aimed at boosting anti-tumour cytotoxic T lymphocytes (CTL) are showing promise in the clinic. Naturally occurring responses can be initiated during tumour growth to establish immunosurveillance in which a dynamic process of immunoediting can ensue; immunological pressure from anti-tumour CTL balances tumour elimination against the emergence of tumour escape variants with no accompanying net outgrowth of tumour (8). This process is responsible for shaping the immunogenicity of the tumour (9). Breakdown of this equilibrium leading to tumour outgrowth involves multiple factors, including the balance between T cell activatory (TcR engagement and co-stimulation) and inhibitory signals (exhaustion markers and immunosuppressive cytokines), and evasion of the T cell response through downregulation of antigen processing machinery or antigen loss. Therapeutic approaches designed to tip the balance back in favour of tumour elimination by providing activation agonists or blockade of inhibition is an attractive strategy that is currently investigated (reviewed in (10)). FoxP3+ CD4+ regulatory T cells (Treg) are important in establishing an immunosuppressive tumour microenvironment, and their infiltration into tumours is a negative prognostic biomarker (11) and a significant obstacle to successful immunotherapy, correlating with a poorer outcome in clinical trials (reviewed in (12)). Therefore, Treg depletion as a therapeutic option is being pursued in the clinic, based on studies in mice that showed rejection of transplanted tumours following ablation of Treg with anti-CD25 antibodies (13, 14).

One of the most widely used mouse models for preclinical testing of new immunotherapeutic drugs is the transplantable BALB/c derived colorectal tumour CT26 (15, 16). In this model, we have shown that depletion of Tregs induces robust protective anti-tumour immunity that effects tumour rejection in ∼90% of mice, similar to responses observed in other mouse tumour models (13, 17). The CT26- immune mice developed memory CTL responses and were able to reject a second challenge with CT26 as well as tumour lines of different histological origin following recovery of Tregs to normal levels. Anti-tumour responses in these mice are focussed on two epitopes derived from *gp90*; AH1 (SPSYVYHQF, (18)) and GSW11 (GGPESFYCASW, (17)). The anti-GSW11 response is more sensitive to Treg suppression *in vivo*, illustrated by the fact that functional (IFN-γ-producing) anti-GSW11 CTL can only be detected in tumour draining lymph nodes (tdLN) in the absence of Treg whereas anti-AH1 CTL are detected whether or not Treg are present (17). Anti-GSW11 CD8+ T cells deliver the most potent anti-tumour response characterised by their ability to reject tumours expressing very low levels of antigen (13, 17).

To investigate the basis of differential suppression of the GSW11-specific T cell response further, we utilised peptide-specific tetramers, to detect both functional (IFN-γ^+^) and inactivated (IFN-γ^-^) antigen-specific T cells. We show that in Treg replete tumour-bearing mice GSW11-specific T cells made up the majority of CD8+ tumour infiltrating lymphocytes (TIL), but exhibit an exhausted phenotype characterised by PD-1 expression and absence of IFN-γ production upon stimulation. We also found that Treg depletion permitted the proliferation of GSW11-specific T cells with lower TcR/MHC avidity suggesting that this population may be more efficacious at rejecting tumour than their higher avidity counterparts with the same epitope specificity.

## Materials and Methods

### Mice, Antibodies and *In vivo* depletion

BALB/c mice were bred under specific pathogen-free conditions in Southampton. Female or male mice (6-8 weeks old) were used in all experiments and during experimental procedures mice were housed in conventional facilities. Hybridomas secreting CD25 (PC61, rat IgG1) specific mAb have been described previously (13). For depletion, mice received intraperitoneal (i.p.) injection of 1 mg of mAb PC61 in 100 µl on days −3 and −1 prior to tumour challenge.

### Tumour cells and *In vivo* challenge

CT26 tumour cells (American Type Culture Collection; ATCC) were maintained in RPMI (Sigma) supplemented with 10% FCS (Globepharm), 2 mM L-glutamine, penicillin/streptomycin (Sigma), 50 µM 2-mercaptoethanol, 1 mM sodium pyruvate (Gibco-BRL) and 1mM HEPES (PAA) and confirmed to be mycoplasma free. In all experiments, mice were injected subcutaneously (s.c.) with 10^5^ tumour cells in endotoxin-low PBS. For analysis of PD-L1 expression, tumour cells were stained with α-PD-L1 (10F.9G2; Biolegend) prior to s.c. injection and after 14-25 days of tumour growth. All flow cytometry data acquisition was carried out on a FACS Canto II (BD Biosciences) and all data analysed with FlowJo Software (Treestar). Tumour cells were gated on live, single cells and the proportion of PD-L1^+^ cells assessed.

### DNA construct

The H2-D^d^ single chain trimer (SCT) construct incorporating a *gp120* HIV peptide (a kind gift from Dr. Keith Gould) was mutated into the GSW11 peptide via site directed mutagenesis (SDM) PCR using KOD HotStart polymerase (Merck Biosciences) according to manufacturer’s instructions. The transmembrane domain of H2-D^d^ was substituted for a biotinylation site using overlapping extension PCR. In addition, a disulphide trap was incorporated into the construct (19) to tether the GSW11 peptide onto the MHC I binding grove.

### Tetramer generation

Tetramers were produced with the help and advice of the Cancer Research UK/Experimental Cancer Medicine Centre Protein Core Facility (Cancer Sciences Unit, University of Southampton, Southampton, U.K.) with few modifications. The GSW11-SCT construct containing H2-D^d^, β2m and GSW11 peptide was cloned into the pET-3a expression vector (Novagen) and expressed in BL-21 CodonPlus RIPL cells (Stratagene). Concentrated refolded complexes were purified on a HiLoad 26/60 Superdex 200 column (GE Healthcare). Biotinylation was achieved with 50µM d-biotin and 1µg/ml biotin protein ligase (Avidity) at 16°C overnight and then passed through the column a second time. Biotinylated monomers were dialysed and subsequently stored in 16% glycerol in PBS or tetramerised by incubation with 1:4 molar ratio of PE-labelled streptavidin (Thermofisher) at 4°C. Each batch of tetramers were tested for binding against the GSW11-specific T cell hybridoma, CCD2Z. For the analysis of AH1-specific T cells, AH1-specific dextramers were used (Immunodex).

### Isolation and analysis of antigen-specific T cells and Tregs

Spleens, tumour draining lymph nodes and tumours from CT26 challenged mice (Treg depleted or replete) were harvested between days 3-25 and disaggregated. CD8+ T cell responses to CT26 antigens GSW11 and AH1 were assessed using antigen-specific tetramers and the production of IFN-γ following peptide stimulation. CD8+ T cells, APCs and peptides/tumours were cultured together for 4 hours in the presence of brefeldin A (BD biosciences) before being stained for cell surface α-CD8 (63-6.7; BD biosciences), antigen-specific tetramer/dextramer, α-PD-1 (RMPI-30; eBiosciences) and intracellular α-IFN-γ (XMG1.2; BD biosciences) using the Cytofix/Cytoperm kit (BD biosciences) according to manufacturer’s instructions. Cells were enumerated by flow cytometry. Numbers reported are those above the background response of T cells alone, with no peptide stimulation. Single cell CD8+ lymphocytes were gated and assessed for tetramer binding and expression of IFN-γ. For analysis of PD-1, CD8+ and tetramer+ were gated and assessed for PD-1 and IFN-γ expression. Within these gates the T cell receptor clonality of GSW11-specific T cells was assessed using a panel of 15 Vβ-specific antibodies (BD biosciences). First, total CD8+ T cells were purified using a CD8 magnetic isolation negative selection kit (Miltenyi) according to manufacturer’s instructions. Purified CD8s were stained with GSW11-specific tetramer, α-Vβ kit, α-CD8 and analysed by flow cytometry. For the analysis of Tregs, cells were stained for cell surface α-CD4 (RM- 4-5; BD biosciences), α-CD25 (7D4; BD biosciences), α-PD-1 and α-PD-L1 (10F.9G2; Biolegend) and intracellular staining with α-FoxP3 (FJK-16S; eBiosciences).

### Tetramer competition assay

Spleens and tumour draining lymph nodes were pooled from Treg depleted or replete mice. CD8+ T cells were purified from disaggregated tissues using magnetic isolation by negative selection (Miltenyi). Purified CD8+ T cells were incubated with 50 nM of dasatinib (New England Biolabs) to prevent TcR internalisation before staining with α-CD8, α-TCR β-chain (H57-597; Biolegend) and 5 μg of PE-labelled GSW11- specific tetramers. After two washes, cells were incubated with bleached tetramer at varying ratios of the initial PE-labelled tetramers: 2.5μg, 5μg, 10μg or 20μg per test. Bleached tetramers were tested for no/minimal PE-fluorescence before use. The β-chain TCR staining was included to confirm the decreasing levels of PE-staining was due to the fluorescently labelled tetramer being out-competed and not due to TcR internalisation.

### Statistical analysis

Analyses were performed using Prism software (GraphPad, San Diego, CA). The *p* values were calculated using either two way ANOVA with Dunnett’s post-test or two-tailed unpaired t test (\**p* < 0.05; \*\**p* < 0.01; \*\*\**p* < 0.001; \*\*\*\**p* <0.0001).

## Results

### Tumour infiltrating GSW11-specific CD8+ T cells are exhausted in CT26 tumour-bearing mice

Recent studies using mouse models have shown that many tumour infiltrating anti-tumour CD8+ T cells have an exhausted phenotype, characterised by PD-1 expression and a failure to express cytokines IL-2, TNF-α and IFN-γ (20, 21). Little is known about whether, or to what extent, development of this phenotype relates to T cell specificity. Clearly, a relationship exists inasmuch as T cell activation via TcR is a prerequisite for the induction of exhaustion (reviewed in (22)). Given our previous observations of differential suppression of anti-CT26 responses with different specificity (17), we sought to investigate the induction of exhaustion in both AH1 and GSW11-specific CD8+ T cells in CT26 tumour bearing mice using antigen-specific tetramers as specificity probes for T cell populations, independent of their functional phenotype. Due to the poor binding affinity of GSW11 for H2-D^d^ (17) we utilised single chain trimer tetramers with GSW11 tethered to the binding groove using a short linker polypeptide (19), allowing stable expression of GSW11-D^d^ monomers. In CT26 challenged Treg replete mice, GSW11-specific T cells were the most abundant CD8+ T cell population in tumours, making up >50% of all CD8+ T cell infiltration after 14 days of tumour challenge (Figure 1A, B, C). However, similar to the situation seen in many human cancers, these infiltrating CD8+ T cells do not confer protection, and tumours continue to grow in these animals. The population of GSW11-specific T cells was significantly larger than AH1-specific T cells, which constituted a maximum of 20% of infiltrating T cells at later stages of tumour challenge (d17 and d22; Figure 1B, C). Notably, while the two T cell populations (anti-GSW11 and -AH1 CD8+ T cells) made up the majority of tumour infiltrating CD8+ T cells (>60%) at later time points (d14-22), they were in the minority in early anti-tumour responses (d7-10, Figure 1C). Thus, an initial broad-specificity polyclonal TIL response becomes focussed on two *gp90* derived epitopes during tumour growth (Figure 1C). We next investigated the function, after *ex vivo* peptide stimulation, of GSW11- and -AH1 specific CD8+ T cells harvested from tumours during the course of the challenge. The vast majority of GSW11-specific CD8+ T cells were unable to produce IFN-γ, with a decrease in functional T cells as the tumour progressed (Figure 1D, E). Most of the AH1-specific T cells were also non-functional, although, consistent with earlier studies showing their presence in tdLN and spleen (17), there were significantly more functional AH1-specific T cells at d22 compared to GSW11 (around 6% compared to <1% functional; Figure 1D, E). This minor population of functional tumour-specific T cells were, however, unable to control tumour growth.

**Figure 1.**
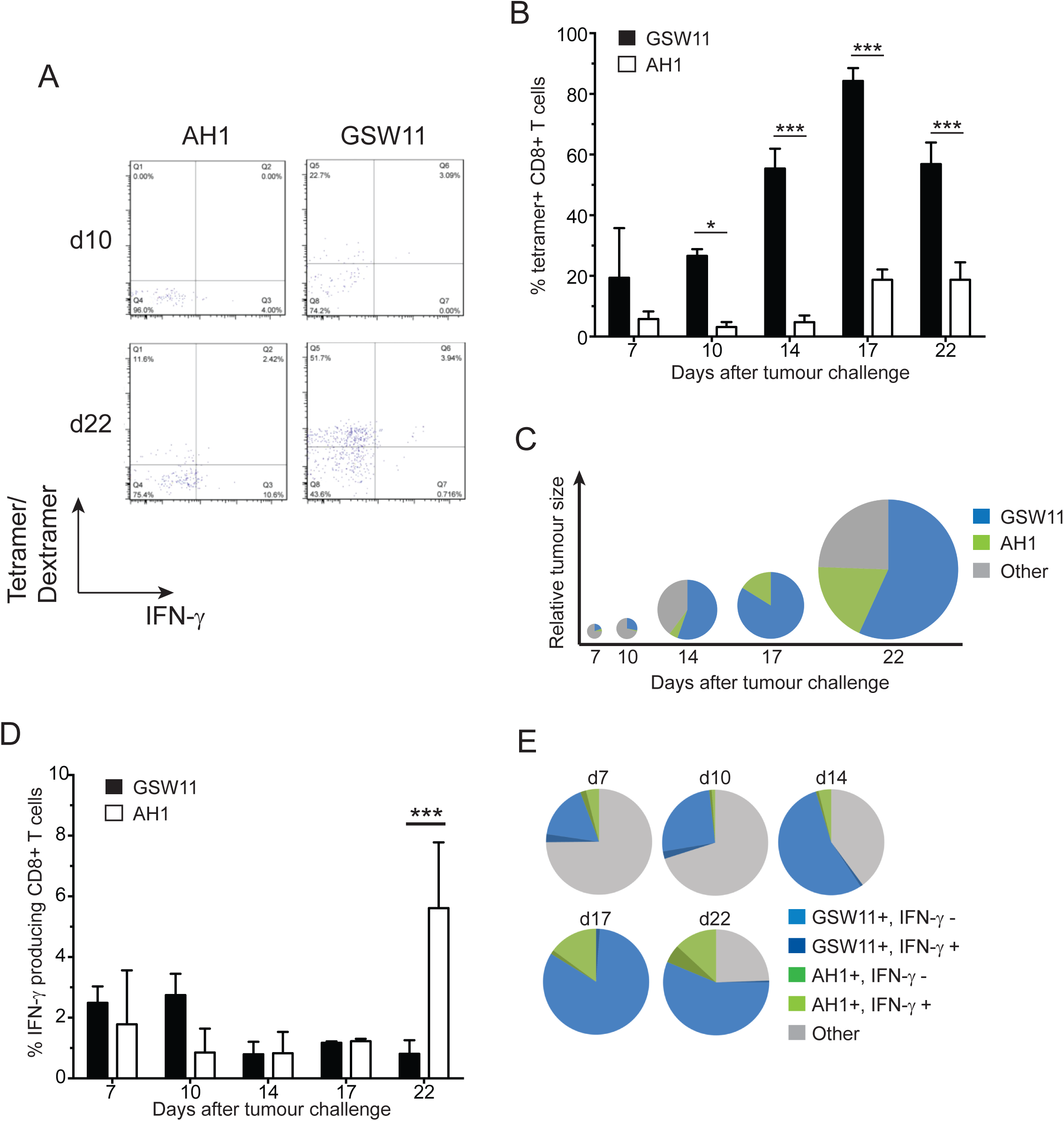
The majority of tumour infiltrating GSW11-specific T cells are non-functional. Balb/c mice were challenged with CT26 tumour cells and the presence of tumour infiltrating AH1- and GSW11-specific T cells was assessed over the indicated time course. (A) Assessment of AH1- and GSW11-specific T cells using tetramer/dextramer and IFN-γ production at d10 and d22. (B) Percentage of antigen-specific T cells detected by tetramer/dextramer. (C) Diagram representing the tumour size (diameter) and relative proportions of antigen-specific tumour infiltrating T cells over time. (D) Percentage of functional AH1- and GSW11-specific CD8+ T cells. (E) Relative proportion of functional and non-functional antigen-specific tumour infiltrating T cells. (B, D; mean and s.e.m. of three mice at each time point; * *p*<0.05, *** *p*<0.001).

To investigate whether the lack of effector function in anti-tumour T cells correlated with the expression of checkpoint molecules commonly associated with T cell dysfunction in tumours, we assessed the expression of PD-1 on GSW11- and AH1- specific T cells following tumour challenge. The majority of non-functional (IFN-γ ^-^) tumour infiltrating T cells of both specificities expressed PD-1 indicating the induction of an exhausted phenotype (50-80%; Figure 2A, B). Anti-GSW11 CD8+T cells acquired the non-functional phenotype more rapidly and to a greater extent compared to AH1-specific T cells. Thus, a greater proportion of GSW11-T cells were PD-1^+^ IFN-γ ^-^ at the first time-point, d7 (*p* =<0.001), which was sustained throughout tumour growth (Figure 2B). Interestingly, a proportion of the small number of functional (IFN-γ^+^) GSW11- and AH1-specific T cells also expressed PD-1, ranging from ∼10% for AH1-specific T cells at day 7 to ∼80% of GSW11-specific T cells at day 22 (Figure 2C), indicating that these T cells are both activated and functional. These results indicate that tumour infiltrating GSW11-specific T cells are highly susceptible to the induction of an exhausted phenotype.

**Figure 2.**
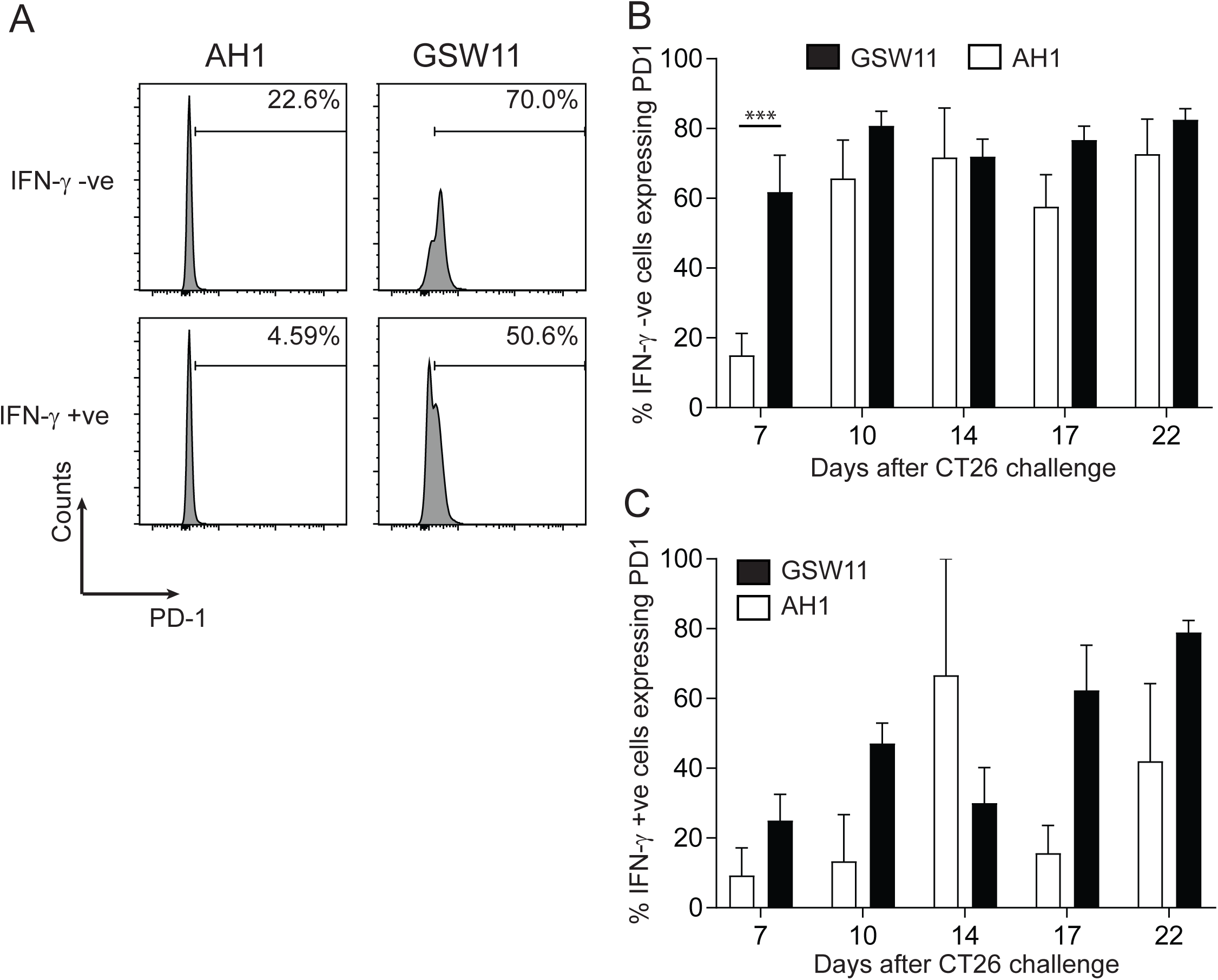
GSW11-specific T cells are exhausted in tumour challenged mice. (A) A representative histogram showing PD-1 expression of functional and non-functional AH1- and GSW11-specific T cells. (B, C) Percentage of PD-1 expressing tumour infiltrating non-functional (B) and functional (C) AH1- and GSW11-specific T cells (B, C; mean and s.e.m. of at least three mice from two independent experiments; *** *p*<0.001).

### The presence of regulatory T cells correlates with PD-1 expression on tumour infiltrating lymphocytes

The tumour microenvironment has been shown to play a crucial role in modulating anti-tumour T cell responses *in vivo* (23, 24) with the presence of tumour-infiltrating Tregs impacting negatively on tumour rejection (11). We therefore investigated the effect of Treg on the induction of exhaustion in infiltrating CD8+ T cells. In a temporal analysis of TIL, Tregs comprised around 15% of total CD4+ T cells at the earliest time point examined (d7). This rose to ∼25% by d10 where it remained for the duration of the tumour challenge (Figure 3A). Interestingly, the accumulation of Tregs in the tumour followed similar kinetics to that observed for the loss of effector function in GSW11-specific T cells (Figure 2B and 1D). Most Tregs expressed PD-L1 in both tumour and the tumour draining LN (tdLN) from the earliest time-point onwards (Figure 3B), suggesting that Treg suppression of GSW11-specific T cells may be mediated via PD-1/PD-L1 interaction. Since tumour-specific T cells also engage with tumour cells for activation, we examined the expression of PD-L1 on tumour cells over 25 days following inoculation. *In vitro* CT26 expressed low levels of PD-L1, however this increased significantly both in terms of the proportion of CT26 expressing, and the level of expression following their seeding and growth *in vivo* (Figure 3C). These results suggest that during initial stages of growth, the tumour provides a pro-inflammatory microenvironment, characterised by IFN-γ producing T cells (and perhaps iNKT/NK cells). This rapidly changes to an immunosuppressive microenvironment in both the tumour and tdLN, characterised by upregulation of inhibitory ligands by the tumour, infiltration of Treg, and upregulation of PD1 on infiltrating tumour-specific CD8+ T cells accompanied by their loss of effector function.

**Figure 3.**
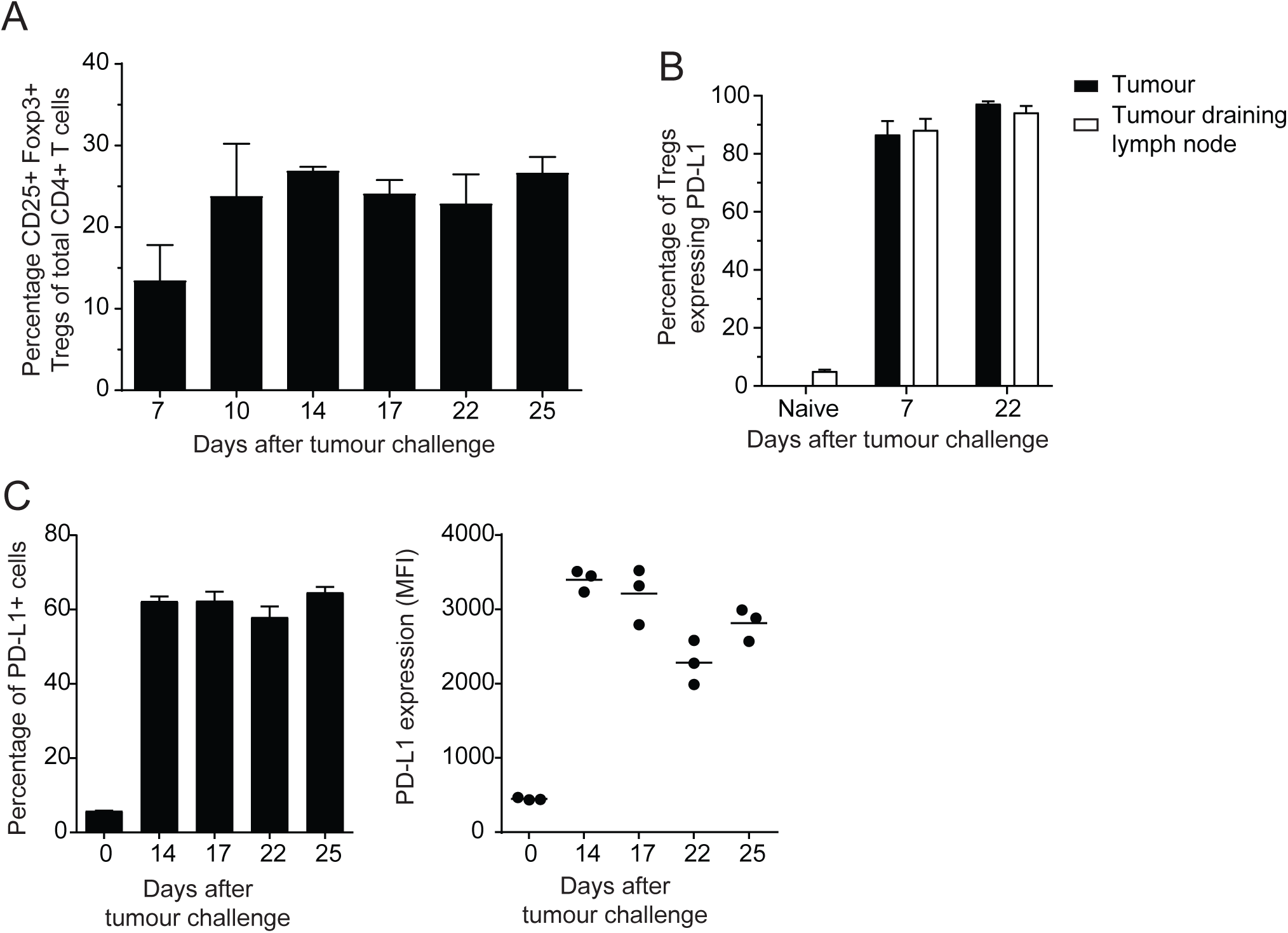
Expression of PD-1 and PD-L1 on tumour infiltrating and LN resident regulatory T cells and tumour cells. (A) Percentage of tumour infiltrating Tregs over time. (B, C) Expression of PD-L1 (B) and PD-1 (C) on Tregs in tumours and tumour draining LN. (D) Percentage and level of PD-L1 expression on CT26 tumour cells prior to and during tumour challenge. (A-C; mean and s.e.m. of three mice at the indicated time points).

### Dysfunction of anti-tumour T cells is induced in the periphery

We next investigated whether GSW11- and AH1-specific T cells are primed effectively to induce T cell effectors. Examination of naïve T cell populations (GSW11 and AH1) showed that both expressed low levels of PD-1 (consistent with previous studies; (25)) with expression on GSW11-specific T cells being slightly greater than AH1-specific T cells (Figure 4A). Following CT26 seeding in Treg replete mice we observed functional (IFN-γ producing) AH1- and GSW11-specific T cells in spleen and tdLN, appearing at d3 and detectable through to the humane end point at d22, with AH1-specific T cells dominating over GSW11-specific T cells at later time points (Figure 4B and C). AH1-specific responses were similar throughout the experiment in lymphoid organs, with the greatest response in spleen (Figure 4C). By contrast, GSW11-specific IFN-γ responses were greatest at early time points (d3- 10) and declined over time (Figure 4B). This confirmed that a co-dominant, functional (IFN-γ) response to AH1 and GSW11 was established in tdLN and spleen immediately following tumour seeding in Treg replete mice and that over time, progressive loss of functional anti-GSW11, but not anti-AH1-specific T cells, was seen. In Treg depleted mice, in which tumours are rejected, GSW11-specific T cells were dominant at most time points, in particular at time points d3-17, in both spleen and tdLN with the greatest response observed in spleen (in line with our previous observations (17) and Figure 4B and C). In addition, the magnitude of anti-GSW11 responses were much greater in Treg depleted compared to Treg-replete mice. AH1-specific responses became co-dominant after d17. Indeed, the magnitude of the anti-AH1 response was not sensitive to the presence or absence of Treg (Figure 4C). Analysis of PD-1 expression on non-functional (IFN-γ ^-^) T cells revealed that a significant proportion (∼20%) of GSW11-specific T cells in the spleen and tdLN expressed PD-1 throughout the time-course, even at the earliest time-point of 3 days; rising to ∼40% at d22 in tdLN (Figure 4D). By contrast, only a very small proportion of IFN-γ ^-^ AH1-specific T cells expressed PD-1, with a maximum of ∼10% of cells in tdLN reached by d22 (Figure 4E). A similar pattern of dysfunctional GSW11- and AH1-specific T cells was observed in the spleens of tumour challenged Treg depleted mice, although, there was a much greater reduction in GSW11-specific T cells compared to AH1, illustrating the differential suppression of anti-CT26 T cells by Tregs ((17) and Figure 4D and E). Importantly, the proportion of dysfunctional GSW11-specific T cells was significantly lower in tdLN of Treg depleted mice (d7 and d22; Figure 4D), suggesting that Treg can exert their effect on T cells in tdLN and spleen and their removal reduces the induction of dysfunction in tumour-specific T cells following priming.

**Figure 4.**
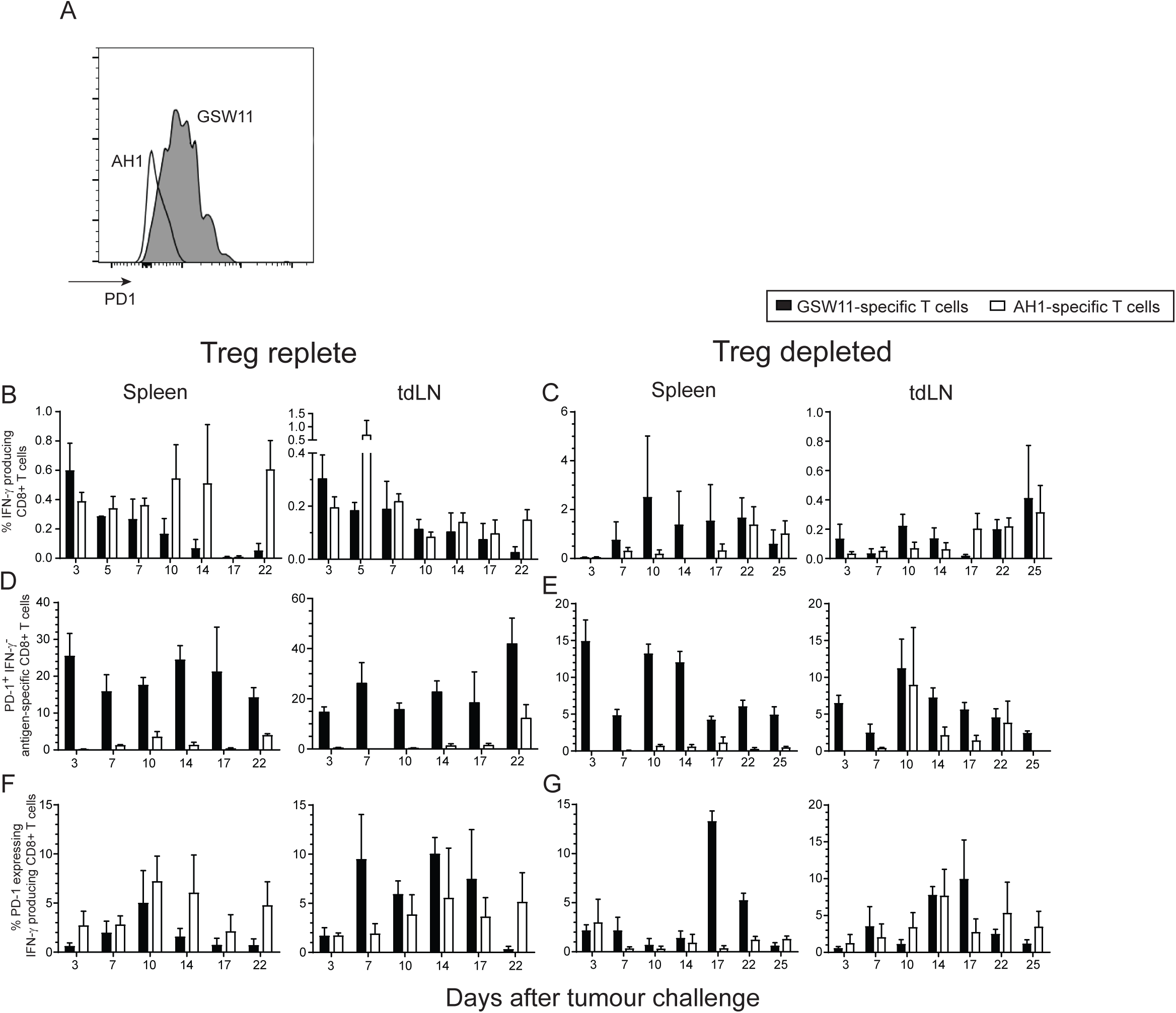
Exhaustion of anti-tumour T cells is induced in the periphery and regulated by regulatory T cells. Treg replete or depleted Balb/c mice were challenged with CT26 and anti-tumour T cell responses in spleens and tumour draining LN assessed. (A) A representative histogram showing the relative level of PD-1 expression on AH1- and GSW11-specific T cells as indicated. (B, C) Percentage of functional GSW11- (B) or AH1- (C) specific T cells in spleen and tdLN in Treg replete and depleted mice following tumour challenge over the indicated time course. (D-G) Proportion of PD-1 expressing non-functional (D, E) and functional (F, G) anti-GSW11 (D, F) or -AH1 (E, G) T cells in spleen and tumour draining LN in Treg replete and depleted mice following tumour challenge. (B-G; mean and s.e.m. of three mice at the indicated time points; d25 data only available for Treg depleted samples).

It is likely that the precursors of IFN-γ ^-^, PD-1^+^ T cells are IFN-γ^+^, PD-1^+^ (22), therefore, to gain a better understanding of the dynamics of transition from functional to dysfunctional phenotypes we assessed PD-1 expression on functional (IFN-γ^+^) CD8+ T cells. In Treg replete mice, levels of PD-1^+^ IFN-γ^+^ GSW11- and AH1- specific T cells were similar, although more AH1-specific T cells were observed at later time points in spleens (d14-22; Figure 4F and G). Similar levels of PD-1^+^ IFN-γ^+^ GSW11- and AH1-specific T cells were observed in Treg depleted mice, however more PD-1^+^ IFN-γ^+^ GSW11-specific T cells were seen at d17 and d22 in spleens (Figure 4F and G). These results indicate that, although the level of primed functional AH1- and GSW11-specific responses is similar, GSW11-specific T cells are more susceptible to the induction of a dysfunctional phenotype.

### Diversity of GSW11-specific T cell responses in Treg depleted mice

Previous studies have shown that the presence of Treg during priming to a transplantation antigen inhibits the priming of T cells with low-avidity TcR (26). To investigate this possibility in the context of Treg-dependent tumour rejection, we first investigated the oligoclonality of the anti-GSW11 T cell response with a view to identifying oligoclonal populations that are preferentially suppressed by Treg. To this end, we determined TcR Vβ usage of GSW11-specific T cells from CT26 challenged Treg replete mice using a panel of V-region specific antibodies. This revealed that the anti-GSW11 response was very diverse with at least 15 different clonotypes observed (Figure 5A and B). Despite the broad response only three Vβ represented >10% of GSW11-specific T cells (Vβ8.1/8.2, Vβ8.3 and Vβ14; Figure 5A) indicating a predominantly oligoclonal response despite the broad Vβ usage. In Treg depleted mice the anti-GSW11 response was similarly broad, although some populations were significantly increased, such as those expressing Vβ3 and Vβ13 and others diminished, such as Vβ10b and Vβ14 compared to Treg replete responses (Figure 5A, B). Intriguingly, some responses such as Vβ8.1/8.2 and Vβ8.3 (which make up ∼30% and ∼12% of the response respectively) were largely unchanged following Treg depletion. These findings suggest that Tregs modulate the anti-tumour GSW11- specific response by preferentially suppressing some T cell clones leading to the expansion of others.

**Figure 5.**
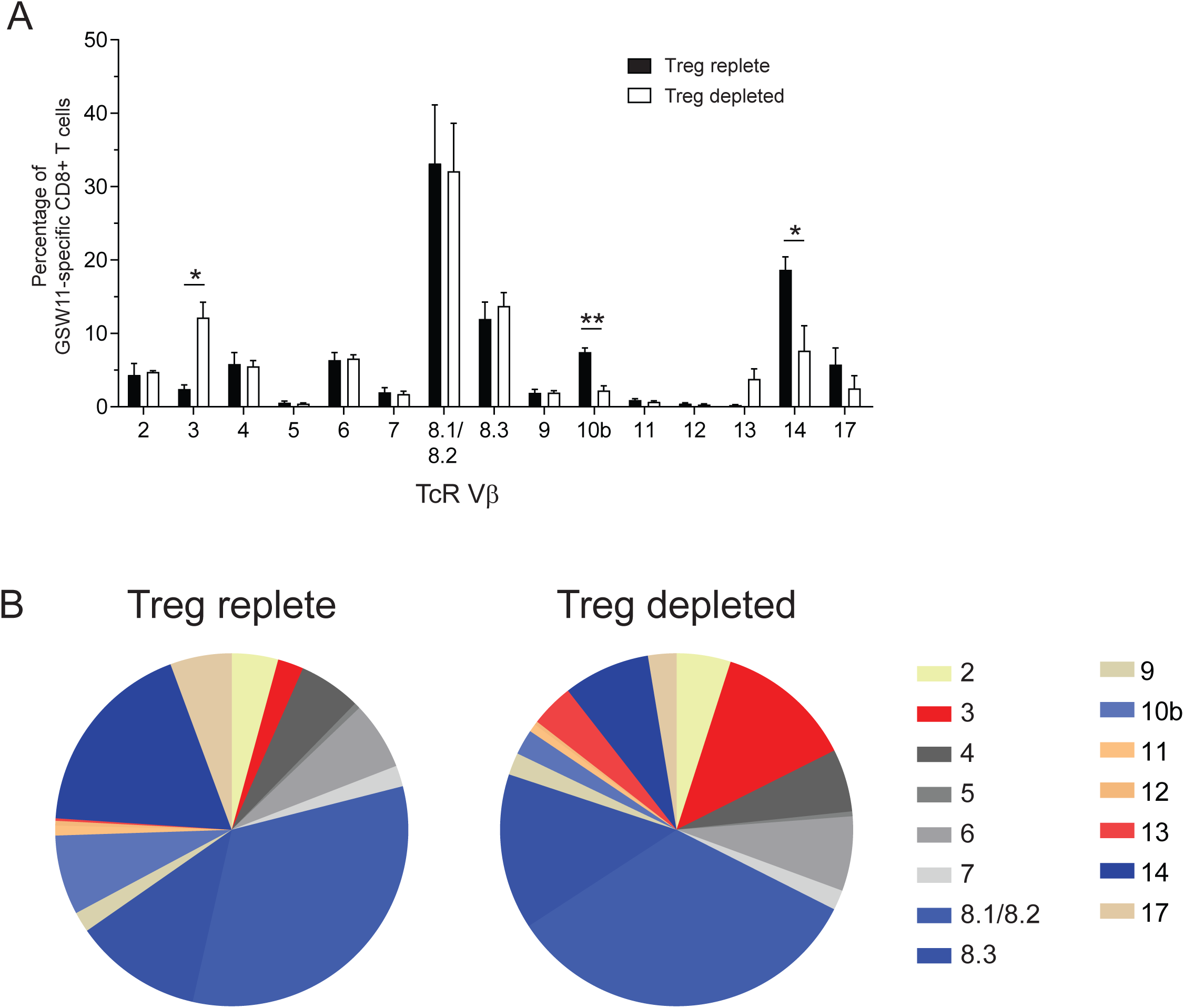
GSW11-specific T cell clonalities are modulated by regulatory T cells. Treg replete or depleted Balb/c mice were challenged with CT26 and the TcR expression of GSW11-specific T cells assessed. (A) Percentage of GSW11-specific CD8+ T cells expressing the indicating TcR Vβ in Treg replete or depleted CT26 challenged mice. (B) The relative proportion of TcR Vβ usage in GSW11-specific T cells following tumour challenge (blue segments indicate TcR with high avidity and red indicate those with low avidity). (A; mean and s.e.m. of ten mice in three pools, B; mean of ten mice from three pools; * *p*<0.05, ** *p*<0.01).

### Tregs target lower avidity anti-GSW11 T cells

We next estimated the TcR avidity of anti-GSW11 T cell oligoclones that were preferentially suppressed by Treg (Vβ3 and Vβ13) compared to oligoclones that were largely unaffected by the presence of Treg (Vβ8.1/8.2 and Vβ8.3), using tetramer competition assays as described previously (Figure 6A and (27)). The two dominant T cell oligoclones that expanded following Treg depletion, Vβ3 and Vβ13, had a lower avidity compared to Vβ8.1/8.2 and Vβ8.3 T cells, which had similar populations whether Tregs were present or not (Figure 6B and C). In addition, two oligoclones, Vβ10b and Vβ14, which were proportionally better represented in Treg replete mice, displayed a high avidity similar to that observed for Vβ8.1/8.2 and Vβ8.3 T cells (Figure 6B and C). Levels of cell surface TcR β chain were similar regardless of the amount of competing tetramer added (Supplementary figure 1). This confirmed that the reduction in tetramer staining was due to competition and not decreased TcR. In pooled samples from groups of 3/4 mice, these low avidity T cells account for ∼16% of the total anti-GSW11 response in Treg depleted animals compared to ∼2.5% in Treg replete animals (red shades; Figure 5B). In addition, high avidity T cells make up >70% and ∼55% of the total anti-GSW11 T cell response in Treg replete and depleted mice respectively (blue shades; Figure 5B). This indicates that sensitivity of a T cell to Treg suppression might be linked to TcR avidity, with lower avidity T cells preferentially targeted. Therefore, in the CT26 model, Treg depletion results in the elaboration of a tumoricidal GSW11-specific T cell response, which appears to correlate with preferential expansion of some low avidity GSW11-specific sub-clones. The response in Treg depleted mice represents a 6-fold increase in low avidity TcR, which make up ∼1/6 of the GSW11-specific response compared to only 1/40 in Treg replete mice.

**Figure 6.**
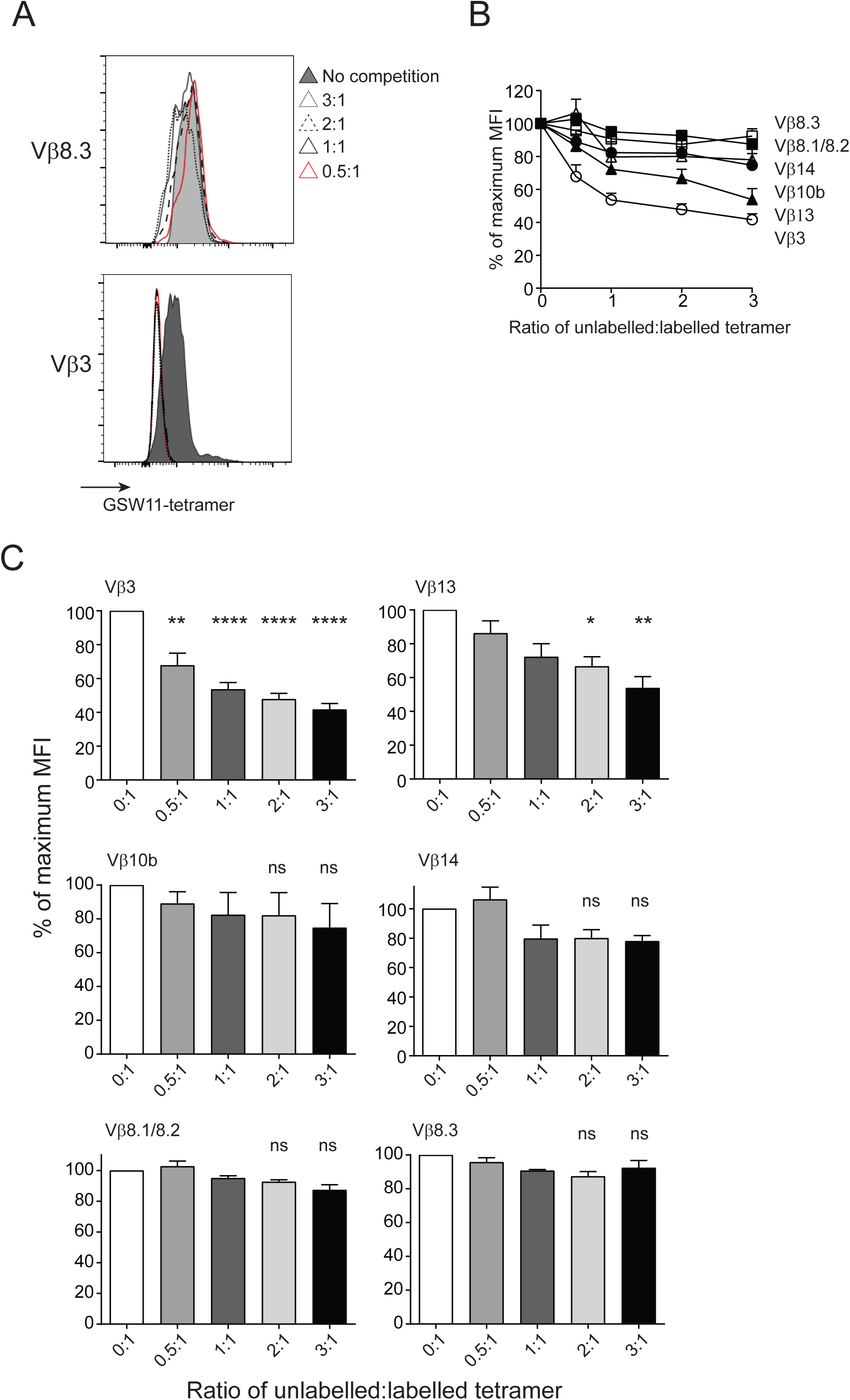
Tumour protective GSW11-specific CD8+ T cells have low TcR avidity. The TcR avidity of anti-GSW11 T cell oligoclones that showed either increased, decreased, or similar levels in tumour challenged Treg depleted mice was assessed using tetramer competition. (A, B) A representative histogram (A) and analysis of relative change in tetramer MFI (B) of the TcR tetramer competition assay in indicated TcR clones. (C) Relative avidity of indicated GSW11-specific T cell oligoclones following tetramer competition (C; mean and s.e.m. of at least three independent experiments; * *p*<0.05, ** *p*<0.01, **** *p*<0.0001).

## Discussion

Immune surveillance of cancers starts with the priming of T cells to tumour associated antigens (TAA) and their infiltration into the diseased tissue. Here, their anti-tumour function is modulated by the evolving microenvironment, often leading to tumour escape via multiple mechanisms including the induction of T cell tolerance/anergy (through lack of co-stimulation during priming) or exhaustion (through the progressive loss of effector function following activation) (22, 28). Using antigen-specific multimers, we show that, in a commercially important preclinical mouse model, growing CT26 tumours are highly infiltrated with tumour-specific CTL recognising one of two non-mutated epitopes (GSW11 and AH1) from a single highly abundant TAA, *gp*90. Most TIL, however, exhibit an exhausted phenotype, which is at least in part mediated by Tregs.

In human cancer, high levels of T cell infiltration generally correlate with good prognosis. This infiltration is marked by a signature that includes increased transcription of genes associated with antigen processing and presentation (MHC I and MHC II), T cell markers (CD8, CD4, CD3) and genes associated with T cell homing (*CCL2*, *CCL3*, *CCL4*, *CXCL9*, *CXCL10 –* the latter two being CD8+T-cell specific), signalling (ICOS, IRF1) and CTL function (granzymes, IFN-γ) (29, 30). These are all consistent with the tumour milieu supporting ongoing peptide:MHC I (pMHC I)-driven T cell proliferation via TcR engagement. However, because of their secretion of the inflammatory cytokine IFN-γ, CD8+ TIL also drive evolution of the immunosuppressive microenvironment including expression of PD-L1, IDO and the infiltration of Tregs (31). In addition, T cells derived from highly infiltrated tumours express the highest levels of inhibitory receptors (such as PD-1) (31). Immunotherapy targets all three key stages of anti-tumour immunity: priming, infiltration and regulation, and the gene signature for T cell inflamed (so-called ‘hot’) tumours encompasses patterns of differentially expressed transcripts associated with positive responses to checkpoint blockade immunotherapy (PD-1, PD-L1, CTLA4) (32).

The CT26 tumour microenvironment resembles cancers with a strong T cell inflamed phenotype, and a signature that includes elevated transcripts for T cell infiltration and activation, CD8, CD4, CD3, CD45, CD62L, CD80, CD86, CD40, OX40L, CD25 and immunosuppression including FoxP3, CTLA-4 and IDO (32). Indeed, we show here that although the tumour mass is infiltrated with tumour-specific CD8+ T cells, they are functionally inert. Despite the obvious differences between spontaneously arising cancer and a transplantable mouse model, studies that have directly compared spontaneous and transplantable tumour models in mice have found only small differences in the tumour-host interaction including immune gene profiles (32, 33). Thus, our time-course of CT26 growth might reasonably approximate to a model for the evolution of the tumour-immune interaction in spontaneously arising cancers in terms of immune-editing and the development of host immune modulation mechanisms within the microenvironment. This is encouraging because transplantable models are more experimentally tractable and permit a more rapid turnaround of preclinical immunotherapy studies.

With this in mind, we observed two key features of the CT26:BALB/c interaction that are relevant to understanding human disease and its response to immunotherapy. Firstly, Treg can differentially suppress different CTL clones recognising TAA and this even applies to CTL recognising the same pMHC I complex; and secondly that differential suppression of GSW11 and AH1-specific T cells, characterised by upregulation of PD-1 expression and loss of effector function (IFN-γ production) was observed very early after tumour challenge (d3) in tdLN. This indicated that a large proportion of GSW11-specific CD8+ T cells were dysfunctional at the site of priming.

The early preferential suppression of anti-GSW11 T cells persists in peripheral tissues where a disparity between dysfunctional GSW11- and AH1-specific T cells is evident in tdLN and spleen. By contrast, in tumours, the difference between GSW11 and AH1 T cells diminishes after d10. This difference between tdLN and tumour indicates that there may be separate mechanisms that operate to establish T cell dysfunction at the early priming and late effector phase. The differential induction of dysfunction in GSW11 at early time points in tdLN, which is significantly reduced in the absence of Tregs, suggests that Tregs play a key role in this process. The rapid expression of PD-L1 on Tregs and PD-1 expression on GSW11-specific T cells indicates this may be an important interaction. Indeed, stimulation of PD-1 (rapidly upregulated following activation) by PD-L1 expressed on antigen presentation cells can cause T cell anergy, defined by reduced proliferative capacity, in a peptide induced model (34). The difference observed in PD-1 expression between AH1- and GSW11-T cells may therefore be a reflection of the relative susceptibility to Treg suppression at the priming stage mediated by PD-1/PD-L1 interaction. Once at the tumour, there is much less difference between AH1 and GSW11 with respect to a dysfunctional ‘exhausted’ phenotype. This change may be due to the increased expression of PD-1 on AH1-T cells, similar to that observed on GSW11-T cells at the tumour, due to continued antigenic stimulation (22). In addition, the more suppressive microenvironment at the tumour site with PD-L1 expression on tumour cells as well as Tregs, which accumulate over time, and the presence of immunosuppressive cytokines such as TGF-β, may overcome any ability of AH1-specific T cells (and to a lesser degree GSW11-T cells) to resist the induction of exhaustion.

The presence of Tregs in several cancer types is a negative prognostic indicator. In addition, tumour-infiltrating Tregs have been shown to express gene signatures, including markers of activation and function, which distinguish them from not only blood Tregs, but also tissue resident Tregs from healthy tissue of the same origin (35, 36). Clinical trials are underway to determine the outcome of agents designed to reduce Treg numbers by targeting the IL-2 receptor or GITR. To date these results have been varied, with daclizumab (anti-CD25) reducing Treg numbers in patients while allowing the induction of T cell responses to targeted tumour antigens (37, 38) and a phase 1 trial of an anti-GITR antibody (TRX518) decreasing Treg numbers in the tumour and circulating blood (39). However, depletion of Tregs in stage IV melanoma with an IL-2/diphtheria toxin conjugate (DAB/IL-2) showed partial responses in only 16.7% of patients, although there was a greater one year survival in partial responders compared to those with progressive disease (80% vs. 23.7%; (40)). This study provides a fuller mechanistic framework for the further rational development of Treg-targeted immunotherapy. Treg suppression of GSW11- specific T cells primarily affected those with lower avidity TcR. These clones expanded in the absence of Tregs, and correlated with survival, suggesting they are important in tumour rejection. It will be interesting to know whether these expanded CTL have a similar profile to the recently described CD103+ CD8+TRM that characterise T cell infiltrates of NSCLC associated with prolonged survival (41). The expansion of protective low Ka T cell clones in Treg depleted mice is consistent with previous studies. For example, Pace et al showed that the presence of Treg during priming to a transplantation antigen increased the affinity of the CD8+ T cell response by inhibiting the priming of T cells bearing low affinity TcR via a mechanism involving CCL3/4 dependent destabilisation of T cell interactions with dendritic cells (26).

The therapeutic efficacy of the low-avidity T cell clones was somewhat unexpected since it is generally assumed that high avidity T cells have a competitive advantage in an immune response due to stronger and prolonged activation signals (42, 43). However, in a situation of persistent antigenic stimulation, as is encountered in the tumour, it is likely that these T cells progress to exhaustion as observed in chronic viral infection. Low avidity CTL may escape the same fate through lower expression of PD-1 or by receiving a TcR signal below the threshold required for exhaustion while maintaining some effector function (44, 45).

The identification of low Ka GSW11-specific TcR, which correlate with protection, may have implications for epitope selection in immunotherapy. Current strategies concentrate on the identification and use of tumour epitopes with a high Ka/slow off-rate in an attempt to induce CD8+ T cell responses with a strong Ka TcR (46-48). This approach has had some success with antigen-specific T cell responses directed to TAA such as NY-ESO-1 and MART-1 and neoantigens. However, only a small proportion of these patients show a partial or complete clinical response (49, 50). These studies show that, while the induction of high-avidity T cells to dominant TAA occur, in a therapeutic setting, it may be more efficacious to induce a broad repertoire of TcR affinities using peptide epitopes presented at sufficient levels regardless of their affinity for MHC. With this in mind, it is notable that the dominant target peptide recognised by TIL in CT26, GSW11, binds weakly to its presenting MHC I, H2-D^d^ with a half-life of ∼20min at the cell surface, though it is presented in high abundance (17). It will therefore be important to understand the relationship between antigen processing/presentation and the induction of low avidity T cells. For example, algorithms predicting the affinity of candidate peptides could be better deployed for selecting candidate epitopes for targeted immunotherapy if used in combination with computational models that take into account antigen abundance and mechanistic details of the antigen processing pathway.

## Supporting information

## Acknowledgements

We thank Leon Douglas and Patrick Duriez from the Cancer Research UK ECMC Protein Core Facility for help and advice with tetramer production and Nasia Kontouli for the purification of the PC61 antibodies.

