## Supplementary material for "Protective low avidity anti-tumour CD8+ T cells are selectively attenuated by regulatory T cells"

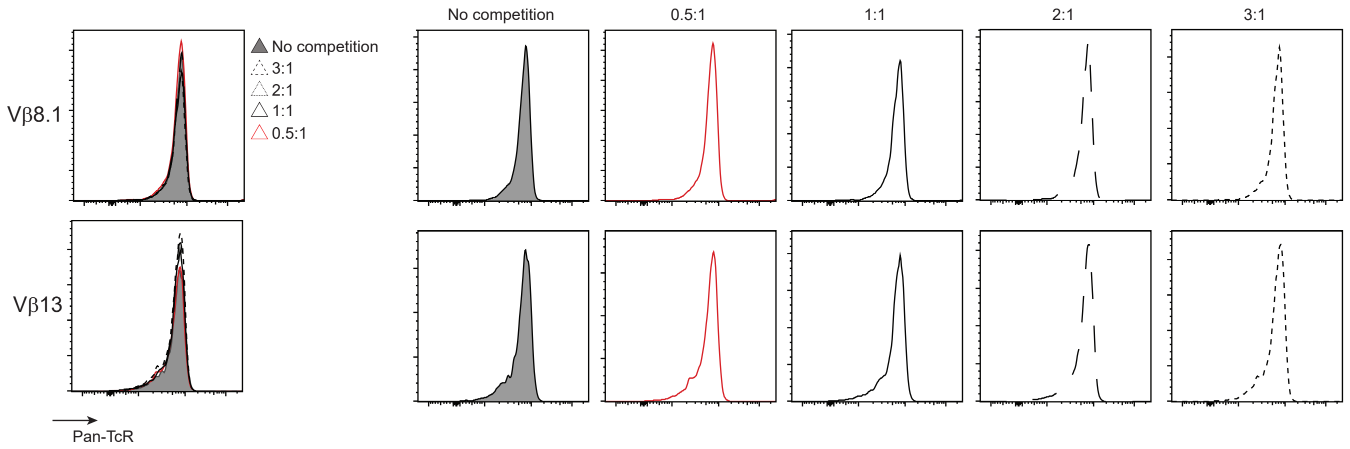


Supplementary figure 1: TcR level is unaffected by tetramer competition
The level of cell surface TcR was assessed during the tetramer competition assay for a high and low avidity TcR. These are representative histograms of three independent experiments.
